# Cytokine profiling identifies circulating IL-2, IL23 and sPD-L1 as prognostic biomarkers for treatment outcomes in Non-Small cell Lung Cancer patients undergoing anti-PD1 therapy

**DOI:** 10.1101/2025.05.13.653410

**Authors:** Kriti Jain, Anika Goel, Deepa Mehra, Deepak Kumar Rathore, Akshay Binayke, Shyam Aggarwal, Surajit Ganguly, Amit Awasthi, Evanka Madan, Nirmal Kumar Ganguly

## Abstract

**Background:** This study investigates the predictive potential of circulating cytokines for response and survival outcomes in patients with advanced non-small cell lung cancer (NSCLC) undergoing immune checkpoint inhibitor (ICI) therapy.

**Materials and Methods:** A cohort of 64 patients with advanced NSCLC receiving ICI therapy were included. Baseline serum samples were collected prior to ICI initiation and profiled using a multiplex cytokine panel. Logistic regression, Cox regression, and Kaplan-Meier survival analysis were employed to assess associations between cytokine levels, therapeutic response, progression-free survival (PFS), and overall survival (OS). Gene expression levels of key cytokines were validated in peripheral blood mononuclear cells (PBMCs) using quantitative real-time PCR.

**Results:** Elevated baseline levels of IL-2, IL-23, and sPD-L1 were significantly associated with clinical response to ICI therapy. Among these, sPD-L1 emerged as an independent predictor of response (AUC = 0.87). Multivariate Cox regression showed IL-2 (HR = 0.67), sPD-L1 (HR = 0.15), and IL-23 (HR = 1.18) were significantly associated with PFS and also predictive of OS. Notably, combined profiling of IL-2 and sPD-L1 enhanced predictive power (AUC = 0.95 for both PFS and OS). RT-PCR analysis of PBMCs corroborated these findings, confirming upregulation of IL-2 in responders and elevated IL-23 expression in non-responders.

**Conclusion:** Baseline cytokine profiling particularly of IL-2, sPD-L1, and IL-23 provides important prognostic and predictive information in advanced NSCLC patients undergoing ICI therapy. These biomarkers may facilitate more personalized approaches to immunotherapy and guide clinical decision-making.

## Introduction

Lung cancer remains one of the leading causes of cancer-related mortality worldwide, affecting both smokers and non-smokers. In India, it is currently ranked as the fourth most common cause of cancer-related deaths, highlighting its aggressive nature and the urgent need for improved therapeutic strategies [1].

Over the past few years, significant advancements have been made in the treatment of lung cancer, evolving from conventional chemotherapy and radiotherapy to more targeted therapies designed to target specific molecular drivers of tumor progression. While these advancements have improved disease management to some extent, their overall effectiveness remains limited, often accompanied by severe side effects and minimal impact on long-term survival [2].

In recent years, immune checkpoint inhibitors (ICIs) therapy has emerged as a groundbreaking approach in lung cancer treatment. These therapies function by stimulating the host immune response, targeting inhibitory pathways such as programmed death-ligand 1 (PD-L1), programmed death-1 (PD-1), and cytotoxic T-lymphocyte-associated protein 4 (CTLA-4), which are exploited by tumor cells to evade immune surveillance. Despite their promising potential, the clinical benefits of ICIs are observed in only 15–45% of patients, highlighting the need for robust predictive biomarkers to guide patient selection [3].

Current biomarkers in clinical use such as PD-L1 expression [3], mismatch repair deficiency (MMR) [4], and tumor mutational burden (TMB) [5] are being used, but they have limitations in accurately forecasting treatment response. This limitation has prompted growing interest in the role of cytokines as potential biomarkers and immunomodulatory agents in lung cancer [6].

Cytokines are essential molecular messengers that regulate immune system communication, enabling a coordinated response against target antigens. While immune signaling often occurs through direct cell-to-cell interaction, cytokines allow for rapid and efficient immune modulation. Their role in cancer treatment has gained significant attention, as they enhance immune cell activation, tumor recognition, and anti-tumor responses. However, due to their pleiotropic functions, redundancy, and capacity for both immune stimulation and suppression, their role in cancer immunity is complex and context-dependent. Understanding this dual nature is essential to harnessing cytokines as therapeutic tools and predictive markers [7].

Investigating cytokine profiles in the tumor microenvironment may offer valuable insights into patient responsiveness to immunotherapy. Their ability to reflect the dynamic interactions between tumor cells and immune components positions them as promising biomarkers for predicting treatment outcomes and guiding clinical decisions in advanced non-small cell lung cancer (NSCLC) [8].

This study aims to explore the potential of cytokines as immunological biomarkers in lung cancer, with a focus on their role in modulating immune responses and predicting patient benefit from immune checkpoint blockade.

## Materials And Methods

### Patients and Treatment

This study included a total of 64 patients with histologically confirmed metastatic solid tumours, including non-small cell lung cancer (NSCLC), melanoma, hepatocellular carcinoma, and head and neck squamous cell carcinoma (HNSCC). All patients were above 18 years of age and were recruited from Sir Ganga Ram Hospital (SGRH), New Delhi between 2019 and 2021. Patients received immune checkpoint inhibitor therapy with Nivolumab. The study was conducted following approval from the Institutional Ethics Committee at Sir Ganga Ram Hospital (Reference No. EC/04/19/1499). Detailed Inclusion and exclusion criteria to enroll patients is as mentioned in **Table S1**. In addition to the patient cohort, 30 healthy individuals who volunteered to be a part of the study were also enrolled as the control group, serving as age- and sex-matched healthy controls. All patients received standard-of-care immune checkpoint inhibitors as follows: Nivolumab: 200 mg intravenously, administered biweekly.

Peripheral blood samples (∼10 mL) were collected from all patients prior to the initiation of ICI therapy (Baseline) using both plain yellow-top tubes and EDTA tubes. These samples were used for detailed immunophenotyping and multi-omics analyses.

### Response Assessment

All patients recruited in this study were followed until either disease progression or death. Treatment response was assessed both clinically and radiologically. Radiological evaluation of disease progression was measured using magnetic resonance imaging (MRI) and therapeutic response was evaluated according to the Response Evaluation criteria in Solid Tumors 1.1 (RECIST 1.1). Clinical assessment of response to Nivolumab was performed at 8-week intervals. Patients were categorized as responders if they demonstrated complete response, a reduction in tumor size, or stable disease. Non-responders were defined as those who exhibited an increase in tumor size or derived no clinical benefit within six months of treatment initiation.

### Sample Collection

#### Blood sample processing-isolation of serum/plasma/PBMCs

##### Isolation of Serum

Blood samples were collected in serum separator tubes (SST) or plain yellow-top vials (BD Vacutainer SST tubes, 367989) and allowed to clot at room temperature for 30 minutes. Following clot formation, samples were centrifuged at 1200 rpm for 10 minutes. The resulting serum was carefully isolated and immediately stored at −80 °C until further use.

##### Isolation of Plasma

Blood samples for plasma collection were drawn into EDTA-containing vial. Samples were centrifuged at 3500 rpm for 15 minutes. The plasma supernatant was collected and stored at −80 °C until analysis.

##### Isolation of Peripheral Blood Mononuclear cells (PBMCs)

For immune monitoring, 8 mL of heparinized blood was collected. The samples were diluted 1:1 with Dulbecco’s Phosphate-Buffered Saline (DPBS) (Sigma Aldrich, D8327) and layered over an equal volume of Histopaque-1077 (Sigma Aldrich, 10771). PBMCs were isolated using a density gradient centrifugation technique at 1200 × g for 35 minutes at room temperature. After centrifugation, red blood cells, granulocytes, and platelets settled at the bottom, while PBMCs were present at the plasma– Histopaque interface. The PBMC layer was carefully aspirated without disturbing the other layers. The collected cells were washed with an equal volume of 1× DPBS and centrifuged at 2000 × g for 10 minutes. A second wash was performed with DPBS, followed by centrifugation at 2000 × g for 7 minutes.

The final PBMC pellet was resuspended in freshly prepared freezing medium consisting of dimethyl sulfoxide (DMSO) (Sigma Aldrich, D8418) and fetal bovine serum (FBS) (Corning, 35-015-CV). Cells were cryopreserved and stored in liquid nitrogen until further use.

### Multiplex cytokine profiling

Stored serum samples isolated from whole blood as per the described protocol, were thawed at room temperature prior to analysis. A panel of 15 cytokines-IL-1α, IL-1β, IL-2, IL-4, IL-10, IL-13, IL-15, IL-17, IL-18, IL-21, IL-23, GM-CSF, TNFα, IFN-γ and sPD-L1 was quantified using the Luminex Discovery Kit (Human premixed multi-analyte kit), R& D Systems, LXSAHM-15 on Luminex Platform (Flex Map 3D, ThermoFisher Scientific, USA).

At first, serum samples were diluted 1:10 using the kit-provided diluent and transferred to polypropylene tubes (ELKAY, 2053-001). Each 96-well plate included a 7-point serially diluted standard curve in duplicate, along with 64 patient samples, 8 of which were run in duplicate. Three batches of Luminex panels were required to analyze 200 patient samples and healthy controls. All procedures were conducted according to the manufacturer’s instructions.

Crystals were completely dissolved by warming the reagents to room temperature. The microparticle cocktail containing all 15 cytokine analytes was prepared by centrifuging at 1,000 × g for 30 seconds, vortexing, and then diluting in the supplied diluent buffer.

Briefly, the assay was performed by adding the microparticle cocktail, diluted serum samples, and cytokine standards to the wells of a 96-well plate, followed by incubation for 2 hours at room temperature on a microplate shaker set to 800 rpm. After incubation, the plates were washed and a biotin-conjugated antibody cocktail was added and incubated for 1 hour. Streptavidin-phycoerythrin (PE) was subsequently added, followed by a 30-minute incubation. A final wash was performed, and microparticles were resuspended in wash buffer.

Washing steps were carried out three times using a magnetic microplate washer. Plates were placed on a magnetic base for 1 minute before aspirating the supernatant. After the final wash, 100μL of wash buffer was added to each well, followed by a 2-minute incubation on the shaker (800 ± 50 rpm).

Plates were read within 90 minutes on the Luminex FlexMap 3D analyzer instrument and data were analyzed using xPONENT software. Values falling below the lower limit of quantification were assigned a value of 1/3 of the lower limit of the standard curve [9].

### RNA Isolation and Real Time PCR

Total RNA was isolated from PBMCs of NSCLC patients (at baseline, before the initiation of ICI) using the TRIZOL reagent (Invitrogen, Grand Island, NY) and quantified using a Nanodrop ND-1000 spectrophotometer (Thermo Fischer, USA). cDNA was prepared from one micrograms of RNase-free DNase treated total RNA using first-strand cDNA Synthesis Kit (applied biosystems Thermo), as per manufacturer’s instructions, using random hexamer primers. PCR reactions were carried in Applied Biosystems, Real-Time PCR System (Agilent Technologies Stratagene Mx3005P) using PowerUp SYBR Green PCR Master Mix (Thermo Fisher Scientific, USA). The detail of the primers (sequences and annealing temperatures) used is given in **Table S2**. Thermal profile for the real-time PCR was amplification at 50°C for 2 min followed by 40 cycles at 95°C for 15 sec, 60°C for 30 sec and 72°C for 1 min. Melting curves were generated along with the mean Ct values and confirmed the generation of a specific PCR product. Amplification of GAPDH was used as internal control for normalization. The results were expressed as fold change of control (Untreated samples (18S)) using the 2^-ΔΔCT^ method. Each experiment was done in triplicates and repeated three times. Statistical significance was determined by Student’s t-test analysis (P<0.05) [9].

### Statistics Analysis

Progression-Free Survival (PFS) was defined as the duration from the first day of immunotherapy to the date of disease progression, death, or last follow-up. Patients without progression or death were censored at the time of their last follow-up. Overall Survival (OS) was defined as the time from the first day of immunotherapy to death from any cause, with patients still alive at the last follow-up censored.

Logistic regression analysis was conducted to determine predictive factors associated with objective response. The area under the curve (AUC) of the receiver operating characteristic (ROC) curves was calculated to assess the predictive accuracy for objective response.

Cox regression analysis was performed to identify predictive factors associated with PFS and OS. ROC analysis was subsequently conducted to validate the hazard ratios calculated using Cox regression and to optimize the cutoff values for cytokine levels. Kaplan-Meier survival curves were generated to compare PFS and OS across different cytokine expression groups.

Variables with *p*<0.1 in univariate analysis were included in the multivariate analysis. Two-sided *p*-values < 0.05 were considered statistically significant. All statistical analyses were performed using Python, specifically the lifelines package [10].

## Results

### Clinical and Pathological Characteristics of the Study Cohort

The study cohort comprised of 64 patients with NSCLC undergoing immune checkpoint inhibitor (ICI) therapy, as shown in **Table 1**. The participants ranged in age from 30 to 81 years. The majority were males (65.6%), while 34.3% were females.

**Table 1.**
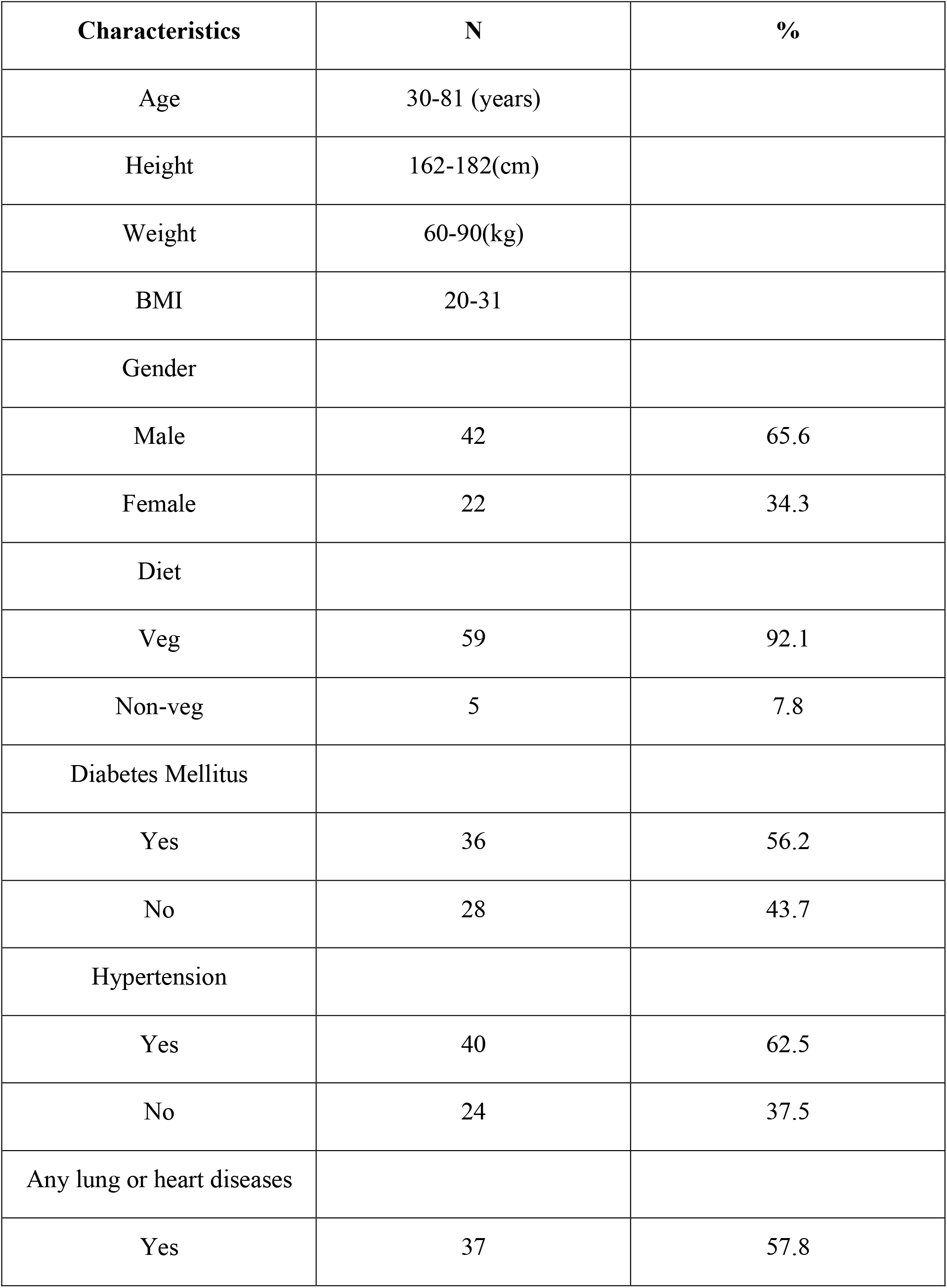
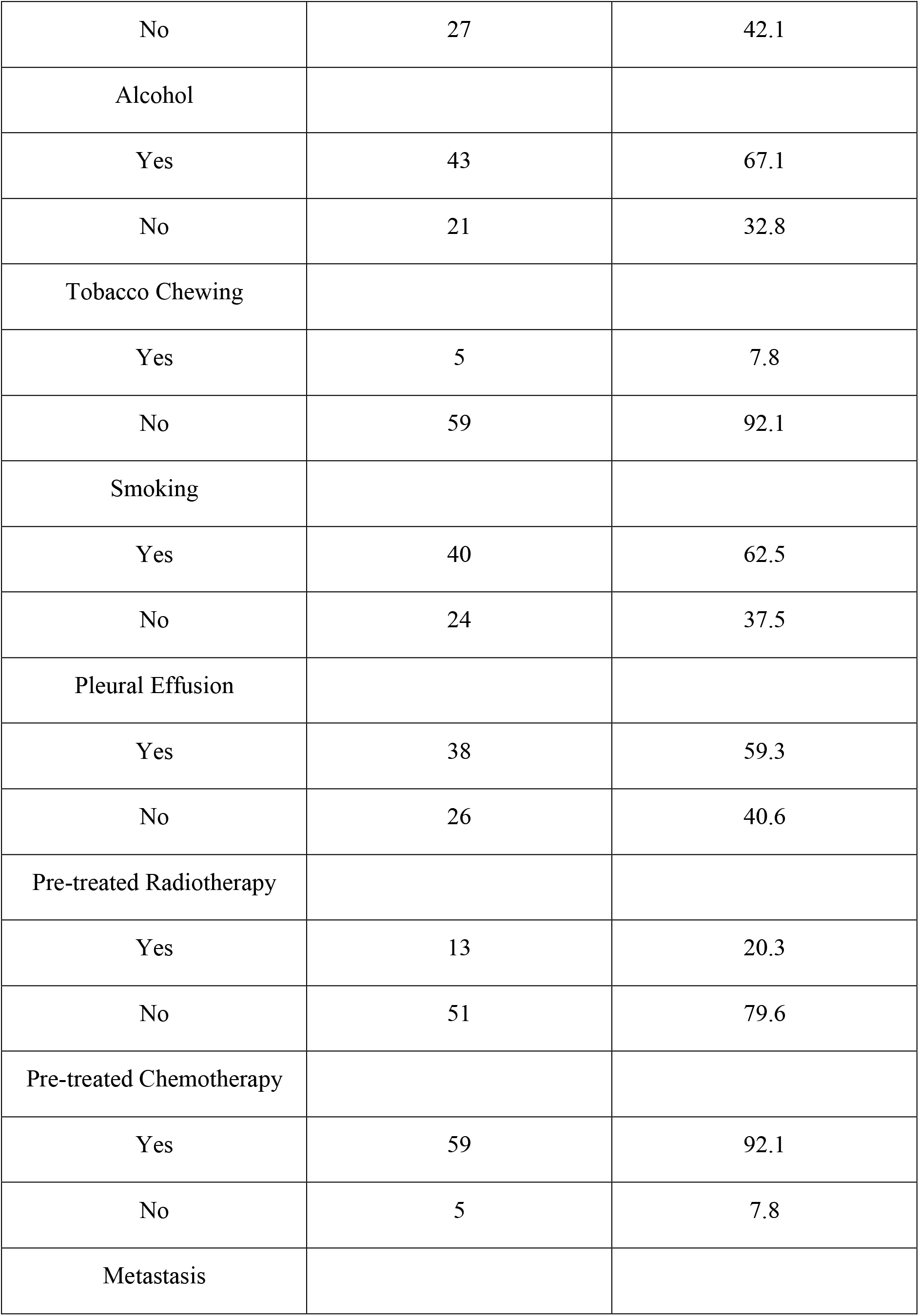
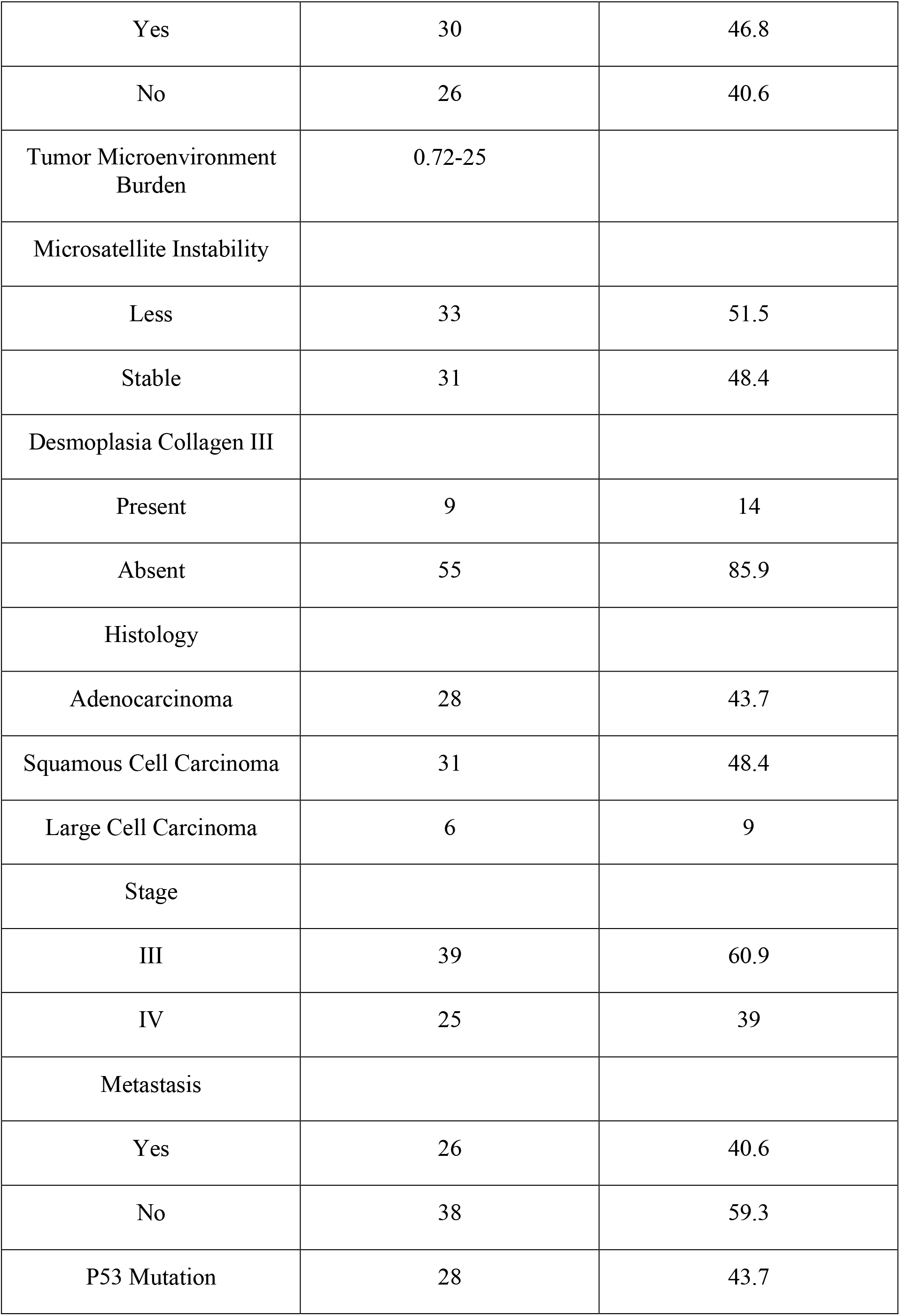
Clinical and demographic characteristics of advanced lung cancer patients (N = 64) The Table presents the clinical and demographic characteristics of 64 patients diagnosed with advanced lung cancer. The data include information on age, height, weight, BMI, gender distribution, dietary habits, comorbid conditions (such as diabetes and hypertension), history of smoking, alcohol consumption, and tobacco use. It also details the presence of pleural effusion, prior treatments (chemotherapy and radiotherapy), metastasis status, tumor microenvironment burden, microsatellite instability, desmoplasia collagen III presence, histological classification, disease staging, and P53 mutation status.

| Characteristics | N | % |
| --- | --- | --- |
| Age | 30-81 (years) |  |
| Height | 162-182(cm) |  |
| Weight | 60-90(kg) |  |
| BMI | 20-31 |  |
| Gender |  |  |
| Male | 42 | 65.6 |
| Female | 22 | 34.3 |
| Diet |  |  |
| Veg | 59 | 92.1 |
| Non-veg | 5 | 7.8 |
| Diabetes Mellitus |  |  |
| Yes | 36 | 56.2 |
| No | 28 | 43.7 |
| Hypertension |  |  |
| Yes | 40 | 62.5 |
| No | 24 | 37.5 |
| Any lung or heart diseases |  |  |
| Yes | 37 | 57.8 |
| No | 27 | 42.1 |
| Alcohol |  |  |
| Yes | 43 | 67.1 |
| No | 21 | 32.8 |
| Tobacco Chewing |  |  |
| Yes | 5 | 7.8 |
| No | 59 | 92.1 |
| Smoking |  |  |
| Yes | 40 | 62.5 |
| No | 24 | 37.5 |
| Pleural Effusion |  |  |
| Yes | 38 | 59.3 |
| No | 26 | 40.6 |
| Pre-treated Radiotherapy |  |  |
| Yes | 13 | 20.3 |
| No | 51 | 79.6 |
| Pre-treated Chemotherapy |  |  |
| Yes | 59 | 92.1 |
| No | 5 | 7.8 |
| Metastasis |  |  |
| Yes | 30 | 46.8 |
| No | 26 | 40.6 |
| Tumor Microenvironment Burden | 0.72-25 |  |
| Microsatellite Instability |  |  |
| Less | 33 | 51.5 |
| Stable | 31 | 48.4 |
| Desmoplasia Collagen III |  |  |
| Present | 9 | 14 |
| Absent | 55 | 85.9 |
| Histology |  |  |
| Adenocarcinoma | 28 | 43.7 |
| Squamous Cell Carcinoma | 31 | 48.4 |
| Large Cell Carcinoma | 6 | 9 |
| Stage |  |  |
| III | 39 | 60.9 |
| IV | 25 | 39 |
| Metastasis |  |  |
| Yes | 26 | 40.6 |
| No | 38 | 59.3 |
| P53 Mutation | 28 | 43.7 |

Among the patients, 62.5% had hypertension, and 57.8% had a history of lung or heart diseases. Additionally, 67.1% of patients reported alcohol consumption, while 62.5% had a history of smoking and 7.8% reported tobacco chewing. In terms of dietary habits, 92.1% of patients followed a vegetarian diet, while 7.8% were non-vegetarians. The presence of pleural effusion was noted in 59.3% of cases.

Prior to ICI therapy, 20.3% of patients had received radiotherapy, and 92.1% had undergone chemotherapy. Metastasis was observed in 46.8% of patients.

Histologically, 43.7% of patients were diagnosed with adenocarcinoma, 48.4% with squamous cell carcinoma, and 9% with large cell carcinoma. Disease staging revealed that 60.9% were at stage III, while 39% were at stage IV. Metastasis was present in 40.6% of cases, and 43.7% of patients exhibited a TP53 mutation.

Tumor microenvironment burden ranged from 0.72 to 25. Microsatellite instability (MSI) analysis showed that 51.5% of patients had low MSI, while 48.4% had stable MSI. Additionally, desmoplasia collagen III was present in 14% of patients, while it was absent in 85.9%.

### Association between cytokine expression with Response to checkpoint inhibitor therapy in NSCLC Patients

To explore circulating non-invasive biomarkers for predicting response to immune checkpoint inhibitor (ICI) therapy in cancer, in depth cytokine profiling was conducted using multiplex Luminex –Flex Map 3D platform. Blood samples were collected at baseline. The panel included 15 cytokines and immune-related factors **i**ncluding interleukins (ILs), interferons (IFNs), tumor necrosis factor (TNF) superfamily members, colony stimulating factors (CSF), chemokines, and growth factors (GFs). These molecules are secreted by immune cells such as monocytes, macrophages, T cells, B cells, and NK cells as well as by certain non-immune cells, including endothelial cells, epidermal cells, and fibroblasts. The cytokines analyzed were: IL-1α, IL-1β, IL-2, IL-4, IL-10, IL-13, IL-15, IL-17, IL-18, IL-21, IL-23, GM-CSF, TNFα, IFN-γ and sPD-L1.

The longitudinal comparison of responders and non-responders was performed using Wilcoxon sign analysis. The results revealed that at the baseline, sPDL-1, IL-2 and IL-23 levels were significantly elevated in responders as compared to non-responders (p<0.05). No statistically significant differences were observed for the remaining cytokines between the two groups **(Figure 1)**.

In concordant with these findings, **Figure 2** presents a heat map comparing the baseline levels of 15 cytokines between responders and non-responders. The data clearly highlight a notable upregulation of soluble PD-L1 (sPD-L1) in responders at baseline.

To validate cytokine expression at the transcript level, RT-PCR was performed on PBMCs isolated from patients with NSCLC. Gene expression analysis revealed distinct transcriptional patterns between responders and non-responders to ICI therapy. Importantly, IL-2 expression was significantly upregulated (∼10 folds) in responders compared to non-responders (p < 0.01), supporting its role as potential predictive biomarker for positive treatment outcomes. In contrast, IL-32, IL-12 and IL-17 showed elevated expression in non-responders (p < 0.01), suggesting a potential association with immune suppression and resistance to ICI therapy as shown in **Figure 3**.

### Association of Cytokines with Response Status in NSCLC Patients undergoing ICI Therapy

To explore the relationship between cytokine levels and treatment response, we performed logistic regression analysis on plasma samples collected prior to immune checkpoint inhibitor (ICI) therapy **(Table S3)**.

Analysis using univariate logistic regression identified TNF-α, PDL-1, IL-10, IL-17, IL-2, IL-13, IL-23 responsive to ICI therapy **(Table S3)**. Further analysis using multivariate logistic regression identified sPD-L1 (Hazard Ratio of 1.51 (95% CI: 1.046-2.81, p <0.005)), as a potential predictive biomarker for response to ICI therapy **(Table 2)**. To assess the predictive performance of cytokine levels, ROC curve analysis was performed. The analysis yielded an area under the curve (AUC) of 0.87 (95% CI: 0.76–0.96) **(Figure 4)**, indicating a high level of diagnostic accuracy for predicting treatment response.

**Table 2.**
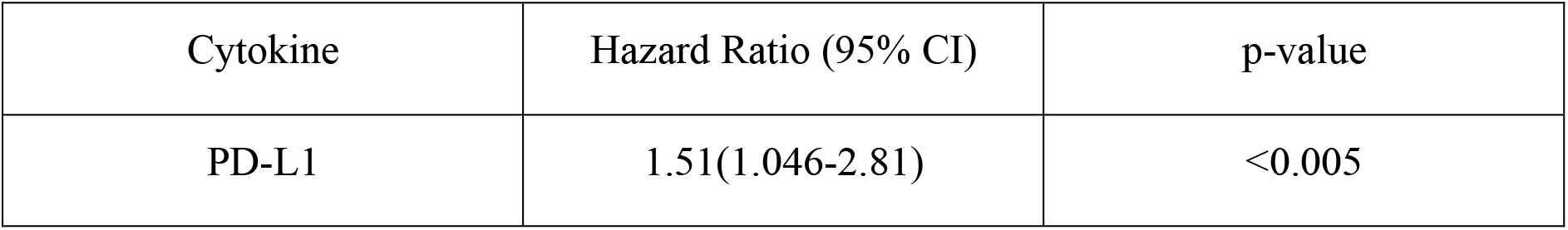
Association between cytokine levels and response status by multivariate analysis. Multivariate logistic regression analysis showing the association between cytokine expression levels and response status. PD-L1 showed a hazard ratio (HR) of 1.51 (95% CI: 1.046-2.81, p <0.005, indicating a strong predictive effect on treatment response.

### Association of Cytokines with Survival Outcomes in NSCLC Patients undergoing ICI Therapy

Univariate Cox regression analysis was performed to evaluate the association of baseline cytokine levels and clinical variables with progression-free survival (PFS) and overall survival (OS) as shown in **Table S4 and Table S5**. Variables demonstrating statistical significance (p < 0.05) in the univariate analysis were subsequently included in the multivariate Cox regression model to identify independent predictors of survival outcomes.

For progression-free survival (PFS), multivariate Cox regression analysis identified IL-2 (HR = 0.67, 95% CI: 0.58–0.79, p = <0.005), sPD-L1 (HR = 0.15, 95% CI: 0.05–0.48, p = <0.005), and IL-23 (HR = 1.18, 95% CI: 1.13–0.98, p = <0.005) as independent predictors of outcome **(Table 3**).

**Table 3.** Association of baseline cytokine levels and clinical variables with progression-free survival (PFS) by multivariate analysis. Multivariate Cox regression analysis showing the association between progression-free survival (PFS) and cytokine expression levels. IL-2 exhibits a hazard ratio (HR) of 0.67 (95% CI: 0.58 – 0.79) with a p-value < 0.005, indicating a significant association with reduced mortality risk. Similarly, sPD-L1 has an HR of 0.15 (95% CI: 0.05 – 0.43) and p-value of <0.005, associated with increased risk of progression. In contrast, IL-23 shows an HR of 1.18 (95% CI: 1.13 – 1.24) with a p-value < 0.005, indicating a significant association with increased mortality risk

| Cytokine | Hazard Ratio (95% CI) | p-value |
| --- | --- | --- |
| IL-2 | 0.67(0.58-0.79) | <0.005 |
| PD-L1 | 0.15(0.05-0.43) | <0.005 |
| IL-23 | 1.18(1.13-1.24) | <0.005 |

Similarly, for overall survival (OS), multivariate analysis revealed that IL-2 (HR = 0.63, 95% CI: 0.54-0.75, p = <0.005), sPD-L1 (HR = 0.29, 95% CI: 0.11–0.80, p = 0.002) and IL-23 (HR = 1.08, 95% CI: –0.1.13, p = <0.005) were significantly associated with survival outcomes **(Table 4)**.

**Table 4.**
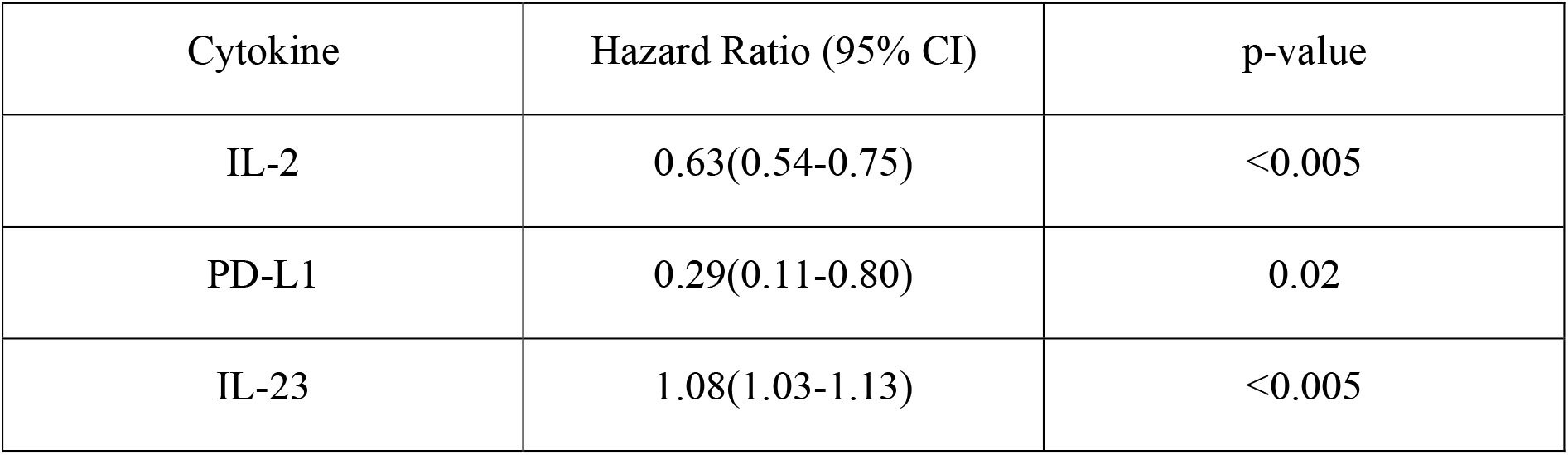
Association of baseline cytokine levels and clinical variables with overall survival (OS) by multivariate analysis. Multivariate Cox regression analysis showing the association between overall survival (OS) and cytokine expression levels. IL-2 exhibits a hazard ratio (HR) of 0.63 (95% CI: 0.54 – 0.75) with a p-value < 0.005, indicating a significant association with reduced mortality risk. sPD-L1 has an HR of 0.29 (95% CI: 0.11 – 0.80) and p-value of 0.02, suggesting a strong protective effect on survival. In contrast, IL-23 shows an HR of 1.08 (95% CI: 1.03 – 1.13) with p-value < 0.005, indicating a significant association with increased mortality risk

Further, Receiver operating characteristic (ROC) analysis was performed for the cytokines identified significant in the multivariate Cox regression. For progression-free survival (PFS,) the area under the curve (AUC) values were 0.92 (95% CI: 0.89 – 0.95) for IL-2, 0.75 (95% CI: 0.69 – 0.81) for sPD-L1, and 0.99 (95% CI: 0.98 – 1.00) for IL-23, indicating strong predictive performance **(Figure 5**). Similar trends were observed for overall survival (OS), with AUC values of 0.92 (95% CI: 0.89 – 0.95) for IL-2, 0.75 (95% CI: 0.69 – 0.81) PD-L1 and 0.99 (95% CI: 0.98 – 1.00) for IL-23 **(Figure 6)**, further supporting their potential as robust prognostic biomarkers.

Kaplan-Meier survival analysis, stratified by cytokine expression levels using ROC-determined cut-offs, further confirmed these associations. For progression-free survival (PFS), higher baseline levels of IL-2 (log-rank p < 0.001) and sPD-L1 (log-rank p < 0.001), along with lower levels of IL-23 (log-rank p < 0.001) were significantly associated with improved outcomes **(Figure 7)**. Similar associations were observed for overall survival (OS), where higher levels of IL-2 (log-rank p < 0.001) and sPD-L1 (log-rank p < 0.001), as well as lower levels of IL-23 (log-rank p < 0.001) were predictive of longer survival **(Figure 8)**.

Combined analysis of IL-2 and PD-L1 revealed complementary predictive power for both progression-free survival (PFS) (HR for IL-2 = 0.86 (95% CI: 0.79 – 0.93) with a p-value = 0.0003, PD-L1 = 0.14 (95% CI: 0.09 – 0.21) with a p-value < 0.0001) and overall survival (OS) (HR for IL-2 = 0.83 (95% CI: 0.76 – 0.91) with a p-value < 0.0001, PD-L1 = 0.15 (95% CI: 0.10 – 0.22) with a p-value < 0.0001). Cox regression analysis confirmed the combined prognostic significance of these cytokines (IL-2, HR= 0.8, PD-L1, HR= 0.15) exhibiting a substantially reduced hazard ratio for both disease progression and mortality **(Table 5 and Table 6)**.

**Table 5.** Multivariate Cox regression analysis considering both IL-2 and PD-L1 together with PFS in relation to survival outcomes. Multivariate Cox regression analysis evaluating the combined impact of IL-2 and PD-L1 expression levels on progression free survival (PFS) in relation to survival outcomes. The hazard ratio (HR) for IL-2 is 0.86 (95% CI: 0.79 – 0.93) with a p-value 0.0003, indicating that higher IL-2 levels are significantly associated with reduced risk of mortality. The HR for PD-L1 is 0.14 (95% CI: 0.09 – 0.21) with a p-value < 0.0001, demonstrating a strong protective effect of higher PD-L1 expression on survival.

| Cytokine | Hazard Ratio (95% CI) | p-value |
| --- | --- | --- |
| Combination |  |  |
| IL-2 | 0.86 (0.79–0.93) | 0.0003 |
| PD-L1 | 0.14 (0.09–0.21) | < 0.0001 |

**Table 6.** Multivariate Cox regression analysis considering both IL-2 and PD-L1 together with OS in relation to survival outcomes. Multivariate Cox regression analysis evaluating the combined impact of IL-2 and PD-L1 expression levels on overall survival (OS) in relation to survival outcomes. The hazard ratio (HR) for IL-2 is 0.83 (95% CI: 0.76 – 0.91) with a p-value < 0.0001, indicating that higher IL-2 levels are significantly associated with reduced risk of mortality. The HR for PD-L1 is 0.15 (95% CI: 0.10 – 0.22) with a p-value < 0.0001, demonstrating a strong protective effect of higher PD-L1 expression on survival.

| Cytokine | Hazard Ratio (95% CI) | p-value |
| --- | --- | --- |
| Combination |  |  |
| IL-2 | 0.83(0.76 – 0.91) | < 0.0001 |
| PD-L1 | 0.15(0.10 – 0.22) | < 0.0001 |

Receiver operating characteristic (ROC) curve analysis of the combined IL-2 and PD-L1 levels further strengthened their predictive power, showing an improved AUC (∼ 0.95) **(Figure 9)** compared to either cytokine alone, demonstrating the enhanced discriminatory capacity of the combined biomarker model for survival prediction.

To further explore the clinical relevance of this combined biomarker model, patients were stratified into four groups based on their IL-2 and sPD-L1 expression levels: Group 1 (high IL-2 / high sPD-L1), Group 2 (low IL-2 / high sPD-L1), Group 3 (high IL-2 / low sPD-L1), and Group 4 (low IL-2 / low sPD-L1). Kaplan–Meier survival analysis demonstrated that patients in Group 1 had significantly improved PFS and OS compared to the other groups (log-rank p < 0.001), highlighting the synergistic effect of IL-2 and sPD-L1 on survival outcomes **(Figure 10)**.

## Conclusion

This study highlights the prognostic and predictive value of circulating cytokines particularly IL-2, sPD-L1, and IL-23 in patients with advanced non-small cell lung cancer (NSCLC) undergoing immune checkpoint inhibitor (ICI) therapy. Elevated baseline levels of IL-2 and sPD-L1 were associated with improved clinical responses and prolonged progression-free and overall survival, while increase IL-23 levels correlated with poorer outcomes, suggesting its potential role in immune suppression and resistance to ICIs. Notably, the combined assessment of IL-2 and sPD-L1 enhanced predictive accuracy, emphasizing the advantage of multi-marker approaches for more precise patient stratification. These findings, validated through both multiplex cytokine analysis and RT-PCR in PBMCs, highlight the value of integrating immune profiling into routine clinical practice to enable more personalized and effective immunotherapy strategies. Future large-scale, multi-center studies are essential to validate these biomarkers and establish standardized protocols for their clinical application.

## Discussion

In this study, we investigated the cytokine profiling of patients with non-small cell lung cancer (NSCLC) undergoing immune checkpoint inhibitor (ICI) therapy, aiming to assess potential biomarkers associated with treatment response and survival outcomes. Our findings reveal distinct cytokine expression patterns between responders and non-responders, suggesting their potential role in predicting treatment outcomes.

A key observation was the differential expression of pro-inflammatory and immunoregulatory cytokines among responders and non-responders. Elevated levels of IL-2 and PDL-1 were significantly associated with favorable clinical responses to ICI therapy, consistent with previous studies linking robust anti-tumor immune response to cytokine-mediated T-cell activation. Conversely, non-responders exhibited increased levels of IL-23 which are known to contribute to an immunosuppressive tumor microenvironment, thereby potentially attenuating the efficacy of ICIs.

IL-2, a well-characterized cytokine has been extensively studied for its dual role in promoting immune responses and maintaining immune tolerance. Recent studies have explored its applications in autoimmune disorders and cancer immunotherapy. Lykhopiy *et al*. (2023) highlighted the role of IL-2 in regulating T cells, particularly its function in modulating regulatory T cells (Tregs) to maintain immune homeostasis in autoimmune conditions such as systemic lupus erythematosus and type 1 diabetes [11]. IL-2 has also been recognized for its ability to stimulate cytotoxic T lymphocytes (CTLs) and natural killer (NK) cells, which are critical in antitumor immunity. Rokade et al. (2024) emphasized the development of IL-2-based therapies that enhance therapeutic efficacy while mitigating toxicity [12]. Similarly, Kim *et al*. (2021) explored how IL-2 and IL-7 support T-cell proliferation and survival, thereby enhancing antitumor responses [13]. Xu *et al*. (2022) found that interventions such as microwave ablation in NSCLC patients can alter systemic cytokine profiles particularly IL-2 and IFN-γ, indicating a shift in immune status post-treatment [14]. Mao *et al*. (2022) further confirmed, through a meta-analysis, that elevated IL-2 levels are associated with improved outcomes in patients receiving ICIs, reinforcing the utility of cytokine monitoring as a predictive tool [15].

Our study also identified high level of sPD-L1 as a significant plasma biomarker, aligning with findings by Shimizu et al. (2023), who reported that sPD-L1 levels dynamically reflect treatment response in patients receiving PD-1 inhibitors [16]. Similarly, Ancel *et al*. (2023) and Machiraju *et al*. (2021) highlighted the role of soluble immune checkpoints and cytokines in predicting ICI efficacy and resistance, further supporting our observations [17-18].

IL-23, in contrast, appears to contribute to immune suppression within the tumor microenvironment. The IL-23/IL-17 axis plays a significant role in immune regulation, particularly in autoimmunity and cancer immunotherapy. Li *et al*. (2024) reported successful management of pre-existing psoriatic arthritis in cancer patients undergoing immune checkpoint inhibitor (ICI) therapy through targeting this axis. This suggests that modulating IL-23 and IL-17 pathways could mitigate immune-related adverse events (irAEs) while preserving anti-tumor immunity [19].

The IL-23/IL-17 axis is increasingly recognized for its role in cancer immunopathology. Wertheimer et al. (2024) demonstrated that IL-23 supports the stability of an effector Treg phenotype within tumors, potentially undermining antitumor immunity [20]. Li *et al*. (2024) further showed that targeting this axis can manage immune-related adverse events in patients undergoing ICI therapy [19]. Liu et al. (2020) underscored the prognostic significance of IL-23 and Th17 cytokines in NSCLC, proposing them as potential markers for disease progression and immune modulation [21].

To further enhance the predictive utility of cytokines, our combined analysis of IL-2 and sPD-L1 demonstrated superior prognostic performance compared to individual markers. Kaplan–Meier analysis of patients stratified by combined cytokine profiles revealed that those with high IL-2 and sPD-L1 levels experienced significantly longer progression-free and overall survival. These findings emphasize the added value of multiplex biomarker models in refining patient stratification and optimizing treatment decisions.

Despite these promising insights, this study has certain limitations. The relatively small sample size and single-institutional design may limit the generalizability of our findings. Additionally, cytokine levels were measured at discrete time points, which may not fully reflect dynamic fluctuations throughout treatment. Future studies involving larger, multi-center cohorts with longitudinal sampling are needed to validate these findings and establish standardized cytokine-based predictive models.

In conclusion, our study underscores the potential utility of cytokine profiling, particularly IL-2, sPD-L1, and IL-23 into clinical practice to improve patient selection and personalize immunotherapy strategies in advanced NSCLC. These biomarkers may not only enhance treatment efficacy but also help anticipate resistance and toxicity. Further research is warranted to refine these biomarkers and explore their applicability in broader patient populations.

## Author Contributions

Ganguly NK conceptualized the whole work and raised funding. Madan E designed experiments, interpreted data and edited the manuscript. Jain K executed the experiments, analyzed the data, performed data interpretation and wrote the manuscript. All patients were treated by Aggarwal S and he provided clinical inputs and interpretation of the clinical data and its correlation. Awasthi A designed experiments and Luminex based work was carried out at his lab. Rathore DK, Binayke A and Mehra D helped in executing Luminex experiments. Ganguly S provided intellectual contribution to this work. All authors listed have made a substantial, direct, and intellectual contribution to the work and approved it for publication.

## Data Availability Statement

The original contributions presented in the study are included in the article/ Supplementary Material. Further inquiries can be directed to the corresponding author.

## Ethics Statement

The study involving human participant was reviewed and approved by SGRH Institutional Ethics Committee (No. EC/04/19/1499). All the patients provided their written informed consent form and willingness to participate in this study and for the publication of any potentially identifiable images or data included in this article.

## Acknowledgments

We would like to acknowledge the patients and their families for cooperation in providing us the samples being used in this study. Also, we would thank Dr. Shrinivas Shinde (Medical Officer-Oncology), Mr. Yogender Tanwar (Research Coordinator) and nursing staff of Medical Oncology Department, Sir Ganga Ram Hospital and Sir Ganga Ram-City Hospital for helping us with blood sample collection. We are also thankful to technical staff of Immunology Lab, Translational Health Science and Technology Institute for providing us the technical support in performing the Luminex based experiments being conducted in this study.

## Supplementary Material

The supplementary material for this article can be found online.

---

**Figure 1.** Cytokine Profiling of Advanced Lung Cancer Patients receiving ICI therapy at Baseline using Luminex The figure represents dot plots showing baseline cytokine levels for various cytokines in responders (green) and non-responders (yellow) to ICI treatment. Each subplot represents a distinct cytokine, with individual data points corresponding to patient measurements. Horizontal lines within each plot indicate the median cytokine levels for each group. Statistically significant differences between groups are denoted by *.

**Figure 2.** Heat map depicting baseline cytokine profiles in responders and non-responders to ICI therapy The heatmap illustrates the expression levels of 15 cytokines at baseline in responders (green) and non-responders (red). The color scale reflects cytokine expression levels, with red indicating higher levels and blue indicating lower levels. The dendrogram on the left clusters cytokines based on similarities in their expression patterns across subjects, highlighting potential clustering of immune response profiles.

**Figure 3.** Cytokine Profiling of Advanced Lung Cancer Patients receiving ICI therapy at Baseline using RT-PCR The figure represents box plots illustrating the comparison of baseline cytokine levels between non-responders (red) and responders (orange) (normalized to Controls) to ICI therapy. Each box represents the interquartile range, with the horizontal line indicating the median value, and the whiskers extending to the minimum and maximum values within 1.5 times the interquartile range. Individual data points are plotted to show variability within each group. Statistically significant differences are marked with asterisks (**, p < 0.01), while “ns” denotes non-significant comparisons.

**Figure 4.** ROC curves illustrating the predictive value of baseline sPD-L1 levels for therapeutic response The ROC curve displays the true positive rate (sensitivity) versus the false positive rate (1 - specificity) across a range of sPD-L1 threshold values. The area under the curve (AUC) is 0.87 (95% CI: 0.76– 0.96), demonstrating strong discriminatory power of sPD-L1 as a biomarker for predicting clinical response to immune checkpoint inhibitor therapy. Higher AUC values indicate greater predictive accuracy.

**Figure 5.** ROC curves illustrating predictive performance of cytokine levels in relation to progression-free survival (PFS): (a) IL-2, (b) IL-23, and (c) PD-L1 Each curve plots the true positive rate against the false positive rate across various thresholds. The area under the curve (AUC) values were 0.92 (95% CI: 0.89–0.95) for IL-2, 0.75 (95% CI: 0.69–0.81) for IL-23, and 0.99 (95% CI: 0.98–1.00) for PD-L1, indicating their respective discriminatory capacities. Higher AUC values reflect stronger predictive power for PFS.

**Figure 6.** Receiver Operating Characteristic (ROC) curves illustrating predictive value of cytokine levels for overall survival (OS): (a) IL-2, (b) IL-23, and (c) PD-L1 Each curve plots the true positive rate against the false positive rate across various thresholds. The area under the curve (AUC) values—0.92 (95% CI: 0.89–0.95) for IL-2, 0.75 (95% CI: 0.69–0.81) for IL-23, and 0.99 (95% CI: 0.98–1.00) for PD-L1—highlight the varying discriminatory capacities of these cytokines to predict OS, with higher AUCs indicating greater predictive accuracy.

**Figure 7.** Kaplan-Meier survival curves for progression free survival (PFS) based on cytokine levels a) IL2 b) PD-L1 and c) IL23, showing stratification into high and low expression groups. Survival differences were assessed using the log-rank test Kaplan-Meier survival curves illustrating progression-free survival (PFS) stratified by cytokine expression levels: (a) IL-2, (b) PD-L1, and (c) IL-23. Each plot categorizes patients into high (orange) and low (blue) expression groups based on cytokine levels, demonstrating distinct survival probabilities over time. The log-rank test p-values for each cytokine indicate highly significant differences in PFS between the groups.

**Figure 8.** Kaplan-Meier survival curves for overall survival (OS) based on cytokine levels a) IL2 b) PD-L1 and c) IL23, showing stratification into high and low expression groups. Survival differences were assessed using the log-rank test Kaplan-Meier survival curves illustrating overall survival (OS) stratified by cytokine expression levels: (a) IL-2, (b) PD-L1, and (c) IL-23. Each plot categorizes patients into high (orange) and low (blue) expression groups based on cytokine levels, demonstrating distinct survival probabilities over time. The log-rank test p-values for each cytokine indicate highly significant differences in PFS between the groups.

**Figure 9.** Receiver Operating Characteristic (ROC) curves illustrating the predictive performance of the combined expression levels of IL-2 and PD-L1 for survival outcomes The left panel represents the ROC curve for predicting progression-free survival (PFS), while the right panel shows the ROC curve for overall survival (OS). Both curves achieve an area under the curve (AUC) of 0.95 (95% CI: 0.89 – 1.00), indicating excellent predictive accuracy. The dashed line represents a random guess (AUC = 0.5), highlighting the superior discriminatory ability of the combined biomarker model.

**Figure 10.** Kaplan-Meier survival curves showing the association of combined PD-L1 and IL-2 expression levels with overall survival (OS) and progression-free survival (PFS). Survival differences were assessed using the log-rank test The left panel illustrates progression-free survival (PFS), and the right panel shows overall survival (OS). Patients were stratified into four groups based on PD-L1 and IL-2 expression: PD-L1^high/IL-2^high (blue), PD-L1^high/IL-2^low (orange), PD-L1^low/IL-2^high (green), and PD-L1^low/IL-2^low (red). Both survival analyses reveal significantly improved outcomes in the PD-L1^high/IL-2^high group. The log-rank test p-values were 4.12 × 10^−28^ for PFS and 2.70 × 10^−29^ for OS, underscoring a strong association between combined biomarker expression and favorable clinical outcomes.

