## Supplementary File for "Cytokine profiling identifies circulating IL-2, IL23 and sPD-L1 as prognostic biomarkers for treatment outcomes in Non-Small cell Lung Cancer patients undergoing anti-PD1 therapy"

**Supplementary Information**

**Table S1: Detailed Inclusion and exclusion criteria of patients enrolled in the study**

| INCLUSION CRITERIA | EXCLUSION CRITERIA |
|---|---|
| Patients with advanced, metastatic/ recurring NSCLC and are eligible for anti- PD-1 immune checkpoint therapy (nivolumab/pembrolizumab) were enrolled in the study. | Patients requiring concurrent anti-cancer therapy during the study period. Patients with communicable diseases (HIV, Hepatitis, etc). |
| Male and female patients in the age group of 18-80 years | Patients who are currently participating or had participated in clinical trials. |
| Patients willing to give informed consent | Patients on wysolove> 10 mg/day |
| Patients who are willing to comply with all study requirements | Patients with brain or subdural metastases are not eligible, unless they have completed local therapy and have discontinued the use of corticosteroids for this indication for at least 4 weeks before starting study treatment. |
| ECOG performance status≤ 2 | Uncontrolled intercurrent illness including, but not limited to ongoing or active infection, symptomatic congestive heart failure, unstable angina pectoris, recent myocardial infarction, cardiac arrhythmia, or psychiatric illness/social situations that would limit compliance with study requirements, or other comorbid condition that investigator believes may compromise participant's condition |
| Women in reproductive age willing to follow adequate barrier contraceptive measures during the conduct of the study. | Patients who have received chemotherapy radiation or biological therapy within 2 weeks, or hormonal therapy within one week before study treatment start, or any investigational drug within 4 weeks before study treatment start. |
| Life expectancy > 4 months | Pregnant and lactating females |

**Table S1.** The table lists the detailed Inclusion and exclusion criteria of patients enrolled in this study.

**Table S2: Primer Sequences for Target Gene Amplification**

| Target gene | | Sequence | Tm ($^{o}$C) |
|---|---|---|---|
| IL-21 | Forward | 5’AGGTCAAGATCGCCACATGA 3’ | 59 |
| IL-21 | Reverse | 5’ TGCTGACTTTAGTTGGGCCT 3’ | 59 |
| PD-L1 | Forward | 5’ CACGGTTCCCAAGGACCTAT 3’ | 59 |
| PD-L1 | Reverse | 5’ GGCCCTCTGTCTGTAGCTAC 3’ | 59 |
| IL10 | Forward | 5’ TTAAGGGTTACCTGGGTTGC 3’ | 62.6 |
| IL10 | Reverse | 5’ TGAGGGTCTTCAGGTTCTCC 3’ | 63.2 |
| IL12 | Forward | 5’ ATGCCCCTGGAGAAATGGTG 3’ | 68.2 |
| IL12 | Reverse | 5’ GGCCAGCATCTCCAAACTCT 3’ | 65.6 |
| TNF-α | Forward | 5’ CCTCTCTCTAATCAGCCCTCTG 3’ | 63.2 |
| TNF-α | Reverse | 5’ GAGGACCTGGGAGTAGATGAG 3’ | 61.7 |
| IL32 | Forward | 5’ AGGCCCGAATGGTAATGCT 3’ | 65.5 |
| IL32 | Reverse | 5’ CCACAGTGTCCTCAG TGTCACA 3’ | 66.5 |

**Table S2:** The table lists the primer sequences used for amplifying specific target genes. Each gene has a designated forward (F) and reverse (R) primer, along with their respective melting temperatures (Tm).

**Table S3: Association between cytokines levels and treatment response by univariate analysis**

| FACTOR | MEDIAN | HAZARD RATIO | P-VALUE |
|---|---|---|---|
| **AGE** | **65** | **1.144218 (1.048212-1.249018)** | **0.002587** |
| SEX | NA | 0.625 (0.22132-1.764981) | 0.374894 |

 3

| | | | |
|---|---|---|---|
| HEIGHT | 168 | 1.037923(0.976085-1.103678) | 0.234975 |
| WEIGHT | 69 | 0.974616 (0.899249-1.056299) | 0.531215 |
| BMI | 25 | 0.85089 (0.69238-1.045688) | 0.124731 |
| DIET | NA | 1.777778(0.276459-11.432067) | 0.544553 |
| DIABETES MELLITUS | NA | 1.615385(0.597068-4.370472) | 0.344972 |
| HYPERTENSION | NA | 1.222222(0.443363-3.369306) | 0.698117 |
| ANY LUNG OR HEART DISEASES | NA | 1.184211(1.184211-3.205844) | 0.739324 |
| SMOKING | NA | 1.842105(0.655374-5.17773) | 0.246621 |
| ALCOHOL | NA | 1.272727(0.444954-3.640455) | 0.652887 |
| TOBACCO CHEWING | NA | 0.5625(0.087473-3.617178) | 0.544553 |
| **WEIGH LOSS** | **NA** | **3.174603(1.114912-9.039374)** | **0.030487** |
| PLEURAL EFFUSION | NA | 1.235294(0.454465-3.35769) | 0.678741 |
| PRE-TREATED RADIOTHERAPY | NA | 1.037037(0.305794-3.516901) | 0.953456 |
| PRE-TREATED CHEMOTHERAPY | NA | 0.738095(0.114817-4.744808) | 0.749061 |
| **HISTOLOGY** | **NA** | **2.955556(1.051512-8.30738)** | **0.039857** |
| STAGE | NA | 0.928571(0.339572-2.539212) | 0.885194 |
| METASTASIS | NA | 0.809524(0.297824-2.200391) | 0.678741 |
| **MICROSATELLITE INSTABILITY** | **NA** | **3.230769(1.156921-9.022108)** | **0.025211** |

| | | | |
|---|---|---|---|
| DESMOPLASIA COLLAGE III | NA | 3.57E+12(0-inf) | 0.99996 |
| **TP53P GENE MUTATION** | **NA** | **2.955556(1.051512-8.30738)** | **0.039857** |
| **TUMOR MICROENVIRONMENT BURDEN** | **8.775** | **1.232367(1.108194-1.370453)** | **0.000115** |
| **TNF-α** | **5.615161058** | **0.92918(0.866243-0.996689)** | **0.040107** |
| **PD-L1** | **0.513403** | **1.458845(1.052671-2.021742)** | **0.023313** |
| **IL-10** | **0.9996894169** | **0.436349(0.220681-0.862788)** | **0.017113** |
| IL-1β | 1.000162318 | 0.837578(0.652729-1.074775) | 0.163574 |
| IFN-ϒ | 0.6862981773 | 0.688172(0.325249-1.456055) | 0.328397 |
| IL-1α | 0.999620582 | 1.004545(0.452977-2.227734) | 0.991096 |
| IL-4 | 0.7833804353 | 0.91285(0.756402-1.101657) | 0.341795 |
| **IL-17** | **0.9998377747** | **17.62385(2.189363-141.867849)** | **0.00701** |
| **IL-2** | **1.000144817** | **18.26174(3.172607-105.115824)** | **0.001143** |
| GMCSF | 1.000296505 | 1.040435(0.878238-1.232587) | 0.646657 |
| **IL-13** | **0.9999981709** | **15.85767(1.34843-186.487872)** | **0.027972** |
| IL-15 | **4.394997553** | 1.01272(0.968247-1.059235) | 0.5812 |
| IL-21 | 1.000109511 | 0.656599(0.279912-1.540202) | 0.333508 |
| **IL-23** | **0.7759003153** | **0.78093(0.614897-0.991794)** | **0.042609** |
| IL-18 | 1.58785315 | 0.792579(0.424369-1.480274) | 0.465787 |

**Table S3:** This table provides the median values, hazards ratios (HR) with 95% confidence intervals (CI), and p-values for each factor. Statistically significant p-values (< 0.05) are highlighted to denote factors potentially influencing Response Status.

**Table S4: Association of baseline cytokine levels and clinical variables with overall survival (OS) using univariate analysis**

| FACTOR | Median | Hazard Ratio | P Value |
|---|---|---|---|
| AGE | 65 | 0.996857(0.868984 - 1.143546) | 0.964151 |
| SEX | NA | 0.982675(0.292016 - 3.30684) | 0.97748 |
| HEIGHT | 168 | 1.020172 (0.895081 -1.162747) | 0.893568 |
| WEIGHT | 69 | 1.007823 (0.99098- 1.129695) | 0.93568 |
| BMI | 25 | 1.112859 (0.807099 - 1.534453) | 0.51413 |
| DIET | NA | 0.264332(0.02714 - 2.574524) | 0.251925 |
| **DIABETES MELLI TUS** | **NA** | **8.578845(3.7844 -1.672298)** | **0.009986** |
| HYPERTENSION | NA | 1.118074(0.321007 - 3.894266) | 0.86085 |
| ANY LUNG OR HEART DISEASES | NA | 0.686727(0.786147 -0.214857) | 0.526135 |
| SMOKING | NA | 2.326875(0.648501- 8.349016) | 0.195124 |
| ALCOHOL | NA | 0.322775(0.069692 -1.494909 ) | 0.148213 |
| TABACCO CHEWI NG | NA | 0.550827 ( 1.494051 -0.068104 ) | 0.576074 |
| WEIGHT LOSS | NA | 1.626689( 0.416023 - 6.360515) | 0.48433 |
| PLEURAL EFFUSI ON | NA | 1.00156(1.353735 - 0.25908) | 0.998197 |

| | | | |
|---|---|---|---|
| PRE-TREATED RADIOTHERAPY | NA | 1.213673(1.353162 -0.380656 ) | 0.743414 |
| PRE-TREATED CHEMOTHERAPY | NA | 0.103418(0.508587 - 0.006431) | 0.109358 |
| HISTOLOGY | NA | 1.24764(1.380406 - 0.39145) | 0.708323 |
| METASTASIS | NA | 0.549133(0.170949 -1.763953) | 0.314062 |
| STAGE | NA | 0.925442(0.842538 -1.016503 ) | 0.105635 |
| TUMOR MICROENVIRONMENT BURDEN | 8.775 | 0.925442 (-0.171336 -0.016369 ) | 0.105635 |
| MICROSATELLITE INSTABILITY | NA | 0.499921 (0.760038 -0.116875 ) | 0.349797 |
| DESMOPLASIA COLLAGEN III | **NA** | **0.191239(0.039525 - 0.925294)** | **0.039735** |
| TP53P GENE MUTATION | NA | 0.319861(0.08327- 1.22867) | 0.096904 |
| **PD-L1** | **0.513403** | **0.7654567(0.45577-1.56789)** | **0.024437** |
| **TNF-α** | **5.615161058** | **1.218083 (1.000422 -1.4831)** | **0.049511** |
| IL-10 | 0.9996894169 | 1.336698(0.493558 - 3.620167) | 0.568075 |
| IL-1 β | 1.000162318 | 1.682801(0.010803 -262.143255) | 0.83987 |
| IFN-ϒ | 0.6862981773 | 1.5979(0.25546 -9.994853) | 0.616336 |
| IL-1α | 0.999620582 | 4.501389(0.703809 -28.789767) | 0.112068 |
| IL-4 | 0.7833804353 | 1.193862(0.963764 - 1.478896) | 0.104785 |
| IL-17 | 0.9998377747 | 1.083477(0.044898 - 26.146236) | 0.960632 |

| | | | |
|---|---|---|---|
| **IL-2** | **1.000144817** | **0.129847(0.0897497 - 0.1864796 )** | **0.022994** |
| GMCSF | 1.000296505 | 1.85518(0.163942 - 20.99331) | 0.617623 |
| IL-13 | 0.9999981709 | 2.828918(0.070843 - 112.96482 6) | 0.580422 |
| IL-15 | 4.394997553 | 1.056368(0.989709 -1.127516 ) | 0.099165 |
| IL-21 | 1.000109511 | 3.049116(0.443462 -20.964862 ) | 0.257073 |
| **IL-23** | **0.7759003153** | **1.238954(1.191397 - 1.29042)** | **0.035139** |
| IL-18 | 1.58785315 | 0.940561(0.407267 - 2.172176) | 0.885902 |

**Table S4:** This table provides the median values, Hazard's ratios (HR) with 95% confidence intervals (CI), and p-values for each factor. Statistically significant p-values (< 0.05) are highlighted to denote factors potentially influencing OS.

**Table S5: Association of baseline cytokine levels and clinical variables with progression-free survival (PFS) by univariate analysis**

| FACTOR | Median | Hazard ratio | *P-value* |
|---|---|---|---|
| AGE | 65 | 0.996857(0.868984 - 1.1 43546) | 0.964151 |
| SEX | NA | 0.982675 (0.292016- 3.3 0684) | 0.97748 |
| HEIGHT | 168 | 1.007823 (0.899098- 1.1 29695) | 0.893568 |
| WEIGHT | 69 | 1.020172 (0.895081- 1.1 62747) | 0.764761 |
| BMI | 25 | 1.112859(0.807099 - 1.5 34453) | 0.51413 |
| DIET | NA | 0.264332(0.02714- 2.57 4524 ) | 0.251925 |

| | | | |
|---|---|---|---|
| **DIABETES MELLITUS** | **NA** | **8.578845(3.7844- 1.672298)** | **0.009986** |
| HYPERTENSION | NA | 1.118074(0.321007 -3.894266 ) | 0.86085 |
| ANY LUNG OR HEART DISEASES | NA | 0.686727(0.786147 - 0.214857 ) | 0.526135 |
| SMOKING | NA | 2.326875(0.648501 - 8.349016 ) | 0.195124 |
| ALCOHOL | NA | 0.322775(0.069692 - 1.494909) | 0.148213 |
| TABACCO CHEWING | NA | 0.550827(0.068104 - 4.455106 ) | 0.576074 |
| WEIGHT LOSS | NA | 1.00156(0.416023 - 6.360515) | 0.48433 |
| PLEURAL EFFUSION | NA | 1.00156(0.25908 - 3.871862) | 0.998197 |
| PRE-TREATED RADIOTHERAPY | NA | 1.213673(1.353162 - 0.380656) | 0.743414 |
| PRE-TREATED CHEMOTHERAPY | NA | 0.103418(0.508587 - 0.006431) | 0.109358 |
| HISTOLOGY | NA | 1.24764(0.39145 -3.976516) | 0.708323 |
| STAGE | NA | 0.549133(0.170949 -1.763953 ) | 0.314062 |
| METASTASIS | NA | 0.549133(0.170949 - 1.763953) | 0.314062 |
| MICROSATELLITE INSTABILITY | NA | 0.499921(0.760038- 0.116875 ) | 0.349797 |

| | | | |
|---|---|---|---|
| DESMOPLASIA COLLAGEN III | **NA** | **0.191239(0.039525 - 0 .925294)** | **0.039735** |
| **TP53P GENE MUTATION** | **NA** | **0.319861(0.08327- 1.22 867 )** | **0.096904** |
| TUMOR MICROENVIRON MENT BURDEN | 8.775 | 0.777607(0.320169 - 1.8 88607) | 0.578506 |
| TNF-α | 5.615161058 | 1.094069(1.000422 - 1. 4831) | 0.509273 |
| **PD-L1** | **0.513403** | **0.669887(0.570562 - 0.7 86503)** | **0.00509273** |
| IL-10 | 0.9996894169 | 1.336698 (0.493558 - 3. 620167) | 0.568075 |
| IL-1β | 1.000162318 | 1.682801(0.01080 -262. 143255) | 0.83987 |
| IFN-ϒ | 0.6862981773 | 1.5979(0.25546 - 9.9948 53) | 0.112068 |
| IL-1α | 0.999620582 | 4.501389(0.703809 -28. 789767) | 0.112068 |
| IL-4 | 0.7833804353 | 1.193862(0.963764 - 1.4 78896) | 0.104785 |
| IL-17 | 0.9998377747 | 1.083477(0.044898 -26. 146236) | 0.960632 |
| **IL-2** | **1.000144817** | **1.174337 (1.119104 -1.2 32295)** | **0.022994** |
| GMCSF | 1.000296505 | 1.85518(0.163942 - 20.9 9331) | 0.617623 |
| IL-13 | 0.9999981709 | 2.828918(0.07084 - 112. 964826) | 0.58042 |

| | | | |
|---|---|---|---|
| **IL-15** | **4.394997553** | **1.056368(0.989709 -1.127516)** | **0.099165** |
| IL-21 | 1.000109511 | 3.049116(0.443462 -20.964862) | 0.257073 |
| **IL-23** | **0.7759003153** | **0.319092(0.107473 - 0.947396)** | **0.045139** |
| IL-18 | 1.58785315 | 0.940561(0.407267 - 2.172176) | 0.885902 |

**Table S5**: This table provides the median values, hazards ratios (HR) with 95% confidence intervals (CI), and p-values for each factor. Statistically significant p-values (< 0.05) are highlighted to identify factors potentially influencing PFS.


11
